# Engineered symbiotic bacteria interfering *Nosema* redox system inhibit microsporidia parasitism in honeybees

**DOI:** 10.1101/2023.01.13.524015

**Authors:** Haoyu Lang, Hao Wang, Haoqing Wang, Xianbing Xie, Xiaosong Hu, Xue Zhang, Hao Zheng

## Abstract

*Nosema ceranae* is an emergent microsporidia parasite of the European honey bee (*Apis mellifera*), which causes serious nosemosis implicated in honeybee colony losses worldwide. *N. ceranae* is an obligate intracellular eukaryotic parasite that mainly invades the midgut of honeybees. Recent studies find that bee gut microbiota is potentially involved in protecting against parasitism. Here, using laboratory-generated bees mono-associated with gut members, we find that *Snodgrassella alvi* inhibited microsporidia proliferation, potentially via the stimulation of host oxidant-mediated immune response. Accordingly, *N. ceranae* employs the thioredoxin and glutathione systems to defend against oxidative stress and maintain a balanced redox equilibrium, which is essential for the infection process. We knocked down the gene expression using nanoparticle-mediated RNA interference, which targets the γ-glutamyl-cysteine synthetase and thioredoxin reductase genes of microsporidia. It significantly reduces the spore load, confirming the importance of the antioxidant mechanism for the intracellular invasion of the *N. ceranae* parasite. Finally, we genetically modified the symbiotic *S. alvi* to deliver dsRNA corresponding to the genes involved in the redox system of the microsporidia. The engineered *S. alvi* induces RNA interference and represses parasite gene expression, thereby inhibits the parasitism by up to 99.8%. Specifically, *N. ceranae* was most suppressed by the recombinant strain corresponding to the glutathione synthetase or by a mixture of bacteria expressing variable dsRNA. Our findings extend our previous understanding of the protection of gut symbionts against *N. ceranae* and provide a symbiont-mediated RNAi system for inhibiting microsporidia infection in honeybees.

## Introduction

Honey bees (*Apis mellifera*) are pollinators with global economic value responsible for pollinating ecologically and agriculturally valuable crops. For the past decade, a phenomenon known as Colony Collapse Disorder has posed a global threat to honeybee health. Recent studies suggest several factors involved in colony decline, such as parasite and pathogen invasion, pesticide use, and environmental stressors. Honeybees are susceptible to a variety of pathogens and pests, including bacteria^1^, fungi^2^, viruses^3^, *Varroa* destructors^4^, and microsporidian parasites^5^.

The microsporidia are obligate intracellular eukaryotic parasites of honeybees and infect the midgut epithelial cells. Honeybees are mainly infected by two species of microsporidia that cause nosemosis, one of the most severe bee diseases worldwide^6^. *Nosema apis* was initially described in European honeybees and was considered the exclusive parasite species causing nosemosis. Later, another species *Nosema ceranae* was discovered in the Asian honeybee, *Apis cerana*, which is presumed to be the original host, and it may transfer to *A. mellifera* during the past decades^7^. It appears that *N. ceranae* displaces *N. apis* in *A. mellifera*, and the prevalence studies found that *N. apis* infections are becoming rarer than *N. ceranae*^8^. *N. ceranae* transmit via the fecal-oral route and the ingestion of spores from the contaminated hive materials^9^. It can suppress the immune defense mechanism of honeybees, ensuring the infection of epithelial cells^10, 11^. The parasitic infection reduces the lifespan and colony populations of *A. mellifera* and affects host physiology and behaviors^12, 13^.

Honeybees rely on innate immunity to defend against infectious agents, which operate through cellular and humoral mechanisms^14^. The humoral immune system consists of antimicrobial peptide production, which provides defense primarily against bacterial pathogens. For intracellular parasites, insects can clear invading parasites by eliciting oxidative stress^15, 16, 17^. The intestinal epithelial and macrophage cells produce reactive oxygen species (ROS)^18^, including superoxide anion (O_2_^−^), hydrogen peroxide (H_2_O_2_), and hydroxyl radical (HO•). While there is no evidence that ROS is effective in clearing microsporidia, the infection of *N. ceranae* may disrupt the oxidative balance of the honeybee gut^19^.

Host ROS production can be modulated by the gut microbiota to eliminate opportunistic pathogens^18^. Although it is unclear whether the microbiota inhibits the parasitism, *N. ceranae* infection perturbs the native gut composition, which may enhance the intensity of the parasitic microsporidia^20^. The honeybee gut microbiota typically contains five core bacterial members^21^. It has been shown that the bee gut bacteria influence bee health by modulating host immune responses. Specifically, *Snodgrassella alvi* and *Lactobacillus apis* protect honeybees from opportunistic bacterial pathogens by inducing host immune response and AMPs production^22, 23^. Furthermore, the native gut bacteria can be engineered to better improve honeybee health^24^. Leonard *et al*. recently genetically modified *S. alvi*, refining a system to induce RNAi within hosts. By expressing dsRNA to interfere gene expression of *Varroa* mite and DWV, the genetically engineered strains repress DWV and *Varroa* infection^25^. This symbiont-mediated RNAi provides a promising strategy for improving bee resistance against stressors.

Here, we investigate the effect of honeybee gut members on the inhibition of *N. ceranae* invasion. Specifically, *S. alvi* upregulated the expression of host genes related to the ROS-associated immune response and significantly repressed the proliferation of *N. ceranae*. Then, we evaluated the role of the antioxidant system of *N. ceranae* in the adaptation and reproduction in the midgut epithelia. We found that *N. ceranae* mainly employed the thioredoxin and glutathione systems to relieve the intense oxidative stress from the host for parasitism. Finally, we constructed recombinant *S. alvi* to continuously produce dsRNA corresponding to the thioredoxin and glutathione system-related genes of *N. ceranae*, significantly inhibiting the *N. ceranae* proliferation in the midgut cells.

## Results

### Gut bacteria aid in the clearance of the pathogenic *N. ceranae*

We first determined whether the core gut members prevent the invasion of *N. ceranae in vivo*. Gnotobiotic bees mono-associated with different gut bacteria, *Bifidobacterium choladohabitans* W8113, *Bombilactobacillus mellis* W8089, *Lactobacillus apis* W8172, *Gilliamella apicola* B14384H2, and *Snodrgrassella alvi* M0351, were generated in the lab. Microbiota-free honeybees were fed with pure cultures of bacterial strains. After allowing the colonization of symbiotic strains in the gut for seven days, each bee individual was manually infected with *N. ceranae* cell suspensions of 10^4^ spores by oral feeding (Fig. 1A). On Day 17, we quantified the absolute abundance of *N. ceranae* spores in the midguts. It showed that the spore load was significantly lowered in bees mono-colonized with *S. alvi*, while bees colonized by other gut members did not show a significant reduction of *N. ceranae* (Fig. 1B).

**Fig. 1.**
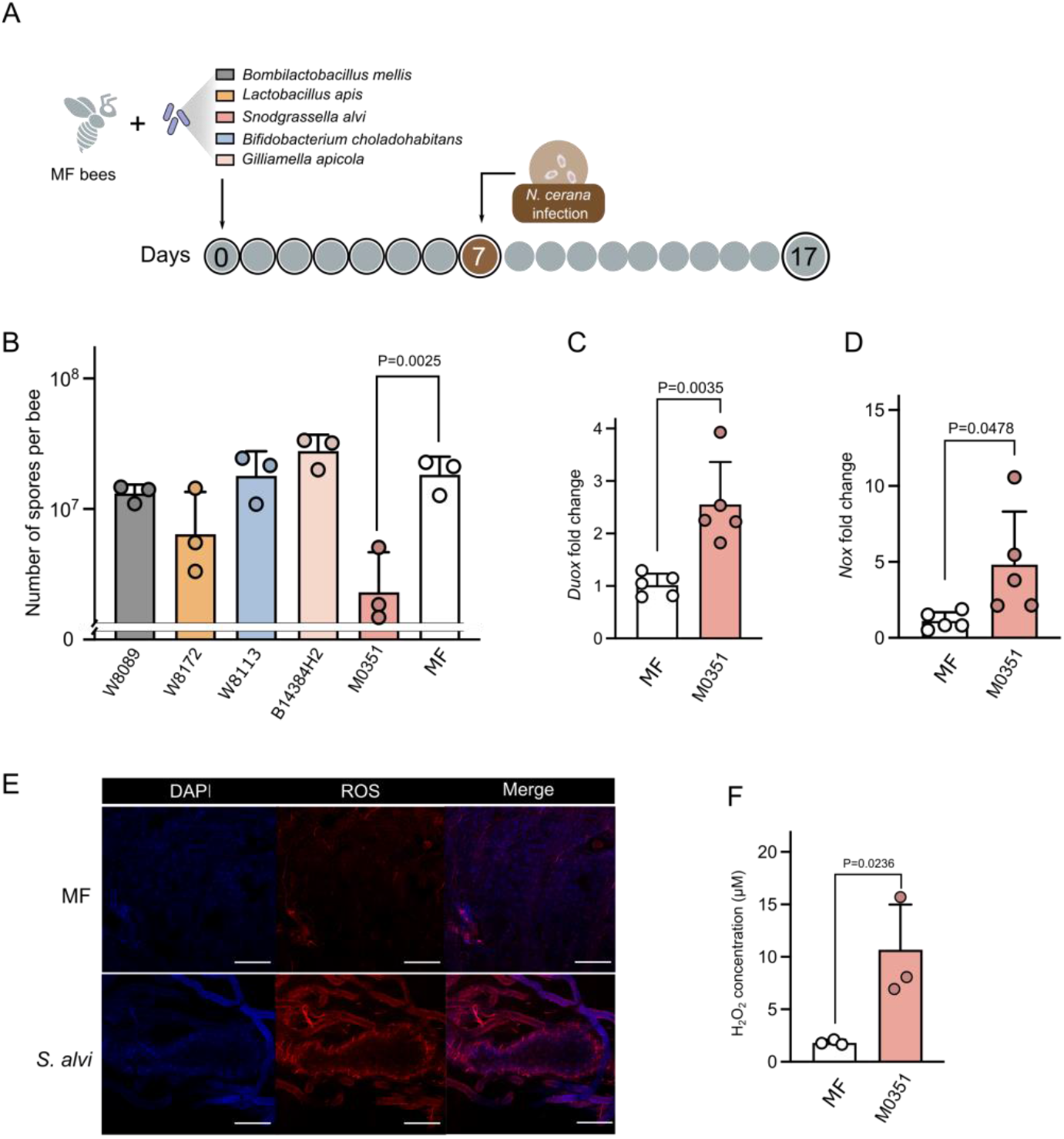
*Snodgrassella* strains protect against *N. ceranae* via the ROS-associated immune response in the honeybee gut. (A) Schematic illustration of experimental design. Microbiota-free (MF) bees were colonized with *B. choladohabitans* W8113, *B. mellis* W8089, *L. apis* W8172, *G. apicola* B14384H2, and *S. alvi* M0351 for seven days and then orally infected with *N. ceranae*. (B) Absolute abundance of *N. ceranae* spores in the midgut 10 days post-infection with *N. ceranae*. (C-D) The expression level of the *Duox* and *Nox* genes in the midgut following *S. alvi* M0351 colonization. (E) Fluorescence staining for ROS signal within the midgut cells of MF and mono-colonized bees with *S. alvi*. (F) H_2_O_2_ concentration in the midgut of MF and mono-colonized bees with *S. alvi*. Scale bars = 250 μm. Statistical analysis was performed by using multiple two-tailed t-test.

Insects can clear parasites from invasion by eliciting oxidative stress, primarily by producing ROS in gut epithelia^26, 27^. Thus, we assessed whether *S. alvi* stimulated the production of ROS in the gut, which may fight against invading intracellular *N. ceranae* parasite. In the honeybee, the production of ROS is mainly regulated by the Nox/Duox NADPH oxidases, as in other insects^28^. We found that the expression of genes encoding Duox and Nox were upregulated in the midgut of bees 24 h post-colonization by *S. alvi* M0351 (Fig. 1C, D). Correspondingly, both the intracellular ROS signal tested by the fluorogenic sensor and the production of hydrogen peroxide (H_2_O_2_) increased in the midgut following the colonization by *S. alvi* (Fig. 2E, F). These results indicate that the colonization of the core gut member, *S. alvi*, triggered the redox response involved in gut immunity, which may inhibit the *N. ceranae* infection in the honeybees.

**Fig. 2.**
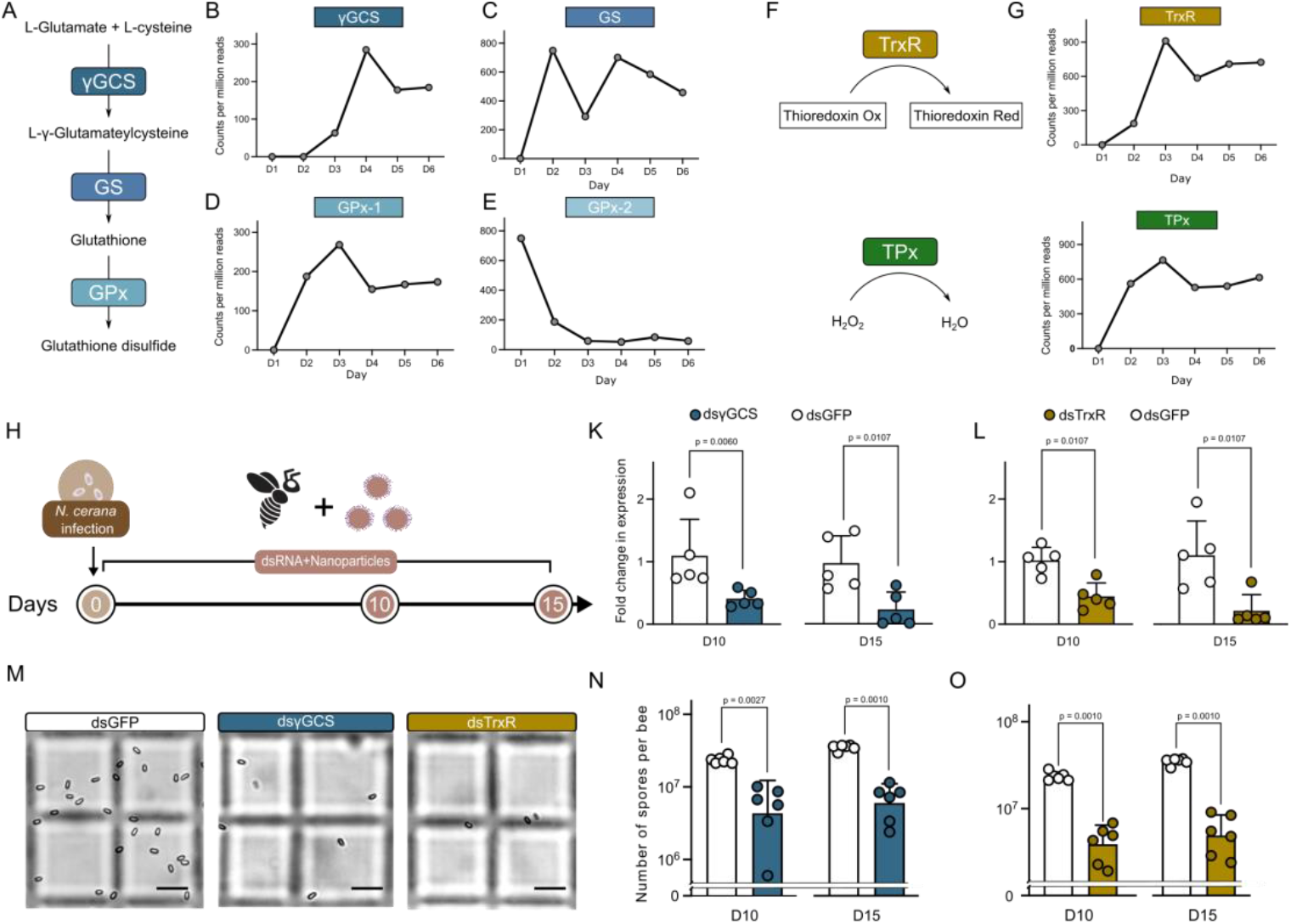
The thioredoxin and glutathione systems of *N. ceranae* are significant for the proliferation in the epithelial cells. (A) The glutathione system forms glutathione by γ-glutamyl-cysteine and glutathione synthetases in *N. ceranae*. (B-E) The expression level of the γGCS, GS, GPx-1, and GPx-2 genes over the infection process. (F) *N. ceranae* possesses a complete thioredoxin system consisting of the thioredoxin reductase and thioredoxin peroxidase. (G) The expression level of the TrxR and TPx genes over the infection process. (H) Knockdown of *N. ceranae* γGCS and TrxR gene expression by feeding nanoparticle-mediated dsRNA. (K-L) Relative expressions of the γGCS and TrxR genes of *N. ceranae* before and after RNAi on Days 10 and 15. (M) The load of *N. ceranae* spores was quantified by microscopy using a hemocytometer. (N-O) Silencing of γGCS and TrxR genes inhibited *Nosema* infection levels. Scale bars = 0.025mm. Statistical analysis was performed by the Mann-Whitney U test.

### *N. ceranae* employs antioxidant systems to adapt and reproduce in the midgut epithelium

We have shown that ROS produced by the bees is implicated in the defense against *N. ceranae*, and typically, the parasites employ endogenous antioxidant systems to relieve intense oxidative stress^29, 30^. To examine the pathways used by *N. ceranae* to resist honeybee gut oxidative stress during infection, we reanalyzed an RNA-seq dataset that documents the changes in gene expression of *N. ceranae* when colonizing the bee gut^31^. *De novo* synthesis of reduced glutathione synthesized by γ-glutamyl-cysteine synthetase (γGCS) and glutathione synthetase (GS) is crucial in the antioxidant defense of *N. ceranae* (Fig. 2A). By following the time-series gene expression profiles, we found that both the expression of γGCS and GS of *N. ceranae* increased along with the infection (Fig. 2B, C). Moreover, glutathione can be further catalyzed by the Glutathione peroxidases (GPx) to reduce H_2_O_2_^32^. We identified that *N. ceranae* possessed two genes encoding GPx in the genome of *N. ceranae*, GPx-1(AAJ76_3500027152) and GPx-2 (AAJ76_3500027978). Interestingly, the expression of GPx-1 increased during the first 3 days of infection, but GPx-2 was downregulated during invasion (Fig. 2D, E). In addition, *N. ceranae* also possesses a complete thioredoxin system, consisting of the key enzymes of thioredoxin reductase (TrxR, AAJ76_5800012528) and thioredoxin peroxidase (TPx, AAJ76_280004776), in defense against oxidative stress (Fig. 2F). We found that the expression of TrxR and TPx genes of *N. ceranae* were elevated from day 2 post-infection (Fig. 2G, H).

To further validate the importance of the thioredoxin and glutathione redox systems for the *N. ceranae* invasion, we knocked down γGCS from the glutathione system and TrxR from the thioredoxin system, respectively. Here, we used the nanoparticle-mediated dsRNA delivery system to improve RNAi efficiency^33^. By feeding the nanoparticle-mediated dsRNA, the mRNA transcript levels of *N. ceranae* γGCS and TrxR genes were reduced by ~80% on Day 10 and Day 15 after inoculation. Microscopic observation confirmed that the proliferation of *N. ceranae* was significantly depressed by both dsγGCS and dsTrxR silencing in the midgut. Altogether, these results indicate that *N. ceranae* probably maintains the redox state by employing the thioredoxin and glutathione systems to relieve the oxidative stress from the host and to adapt and reproduce in the midgut epithelium^34^. This also implies that host ROS-associated immunity is responsible for the defense against intracellular parasitism in honeybees.

### Inhibition of *N. ceranae* infection by engineered *S. alvi*

Since the antioxidant defense is crucial for *N. ceranae* parasitism, we next engineered *S. alvi* strain M0351 to produce dsRNA targeting microsporidian genes. First, we transformed strain M0351 with a stable plasmid pBTK519 expressing GFP from the Bee Microbiome Toolkit platform ^24^ and tested whether it could re-colonize bee gut robustly. The engineered strain M0351 was inoculated into newly emerged bees treated with ampicillin to eliminate native microbiota (Fig. 3A)^25^. We inoculated bee individuals with ~10^5^ CFU of GFP-tagged *S. alvi*. They grew to ~8.0 × 10^7^ CFU/bee after five days of colonization and persisted stably throughout the 15-day experiments (Fig. 3B). The majority of engineered *S. alvi* cells (~80%) remained functional with a high density of fluorescent signal, while some bacterial cells lost the fluorescence in the guts at the endpoint (Day 15; Fig. 3C). While *Snodgrassella* preferentially colonizes the ileum, they also distribute in all compartments of the bee gut^35^. The confocal microscopy showed that 15 days after colonization, the engineered M0351 effectively colonized both the ileum and midguts of 10 inspected bees, showing the same spatial distributions as the wild-type strain (Fig. 3D-F) ^36^. Thus, our results showed that the engineered *S. alvi* could persistently colonize the honeybee ileum and midgut, and the plasmid pBTK519 functioned reliably in strain M0351 throughout the experiments.

**Fig. 3.**
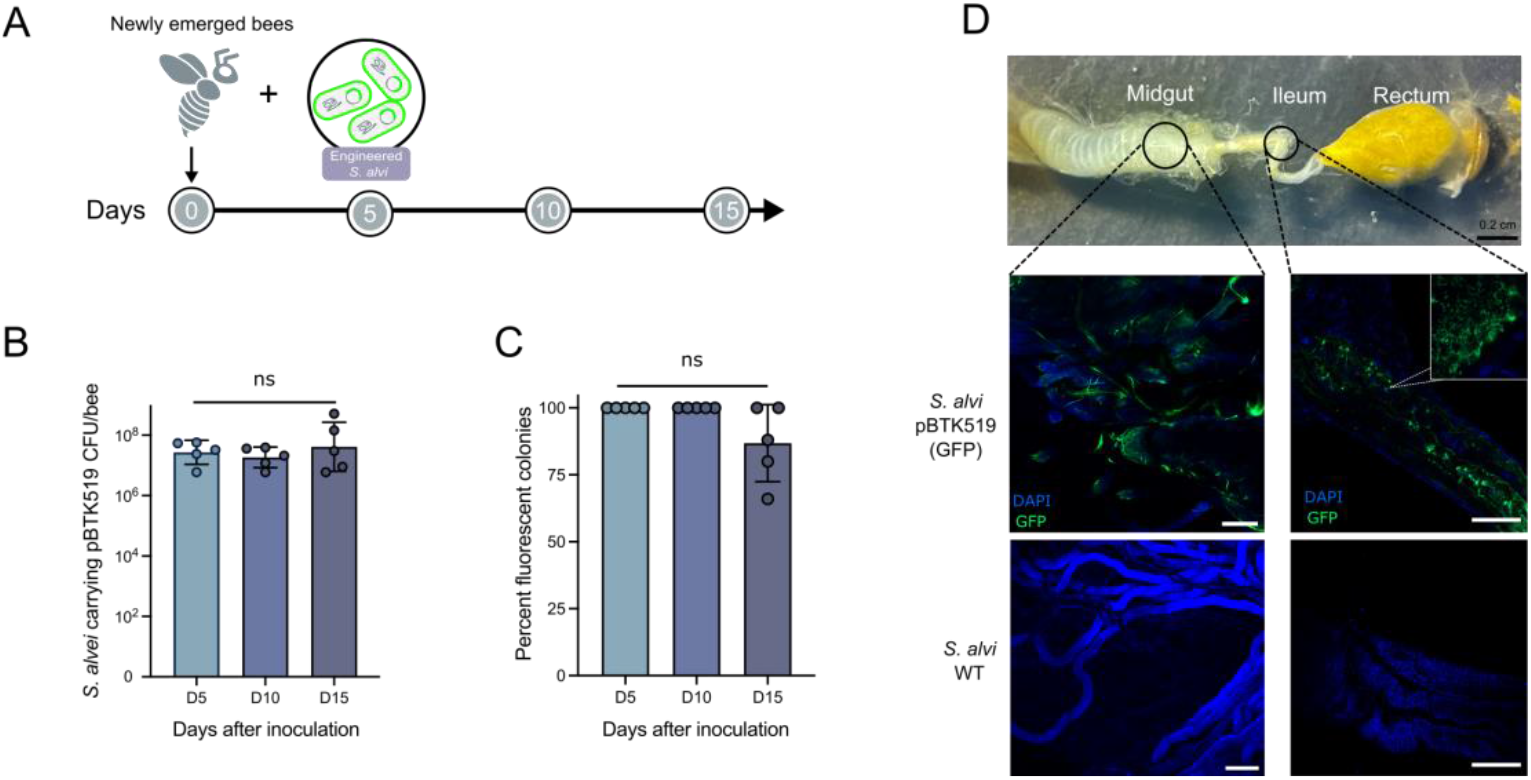
The engineered *S. alvi* M0351 showed stable colonization and function in bee guts. (A) Newly emerged bees were colonized with *S. alvi* transformed with a plasmid expressing a green fluorescent protein. The colonization level was checked on Days 5, 10, and 15. (B-C) Engineered *S. alvi* M0351 stably colonized and expressed GFP continuously over time. Each dot represents an individual bee sample. (D) Engineered *S. alvi* M0351 colonized both the midgut and ileum of bees. Scale bars = 200 μm. Statistical analysis was performed by the Mann-Whitney U test.

We have shown that *S. alvi*-treated honeybees prevent *N. ceranae* infection by triggering ROS production (Fig. 1), and *N. ceranae* employed the thioredoxin and glutathione system to relieve the intense oxidative stress (Fig. 2). Thus, we engineered *S. alvi* M0351 using plasmid pBTK519 to express dsRNA targeting the glutathione and thioredoxin systems of *N. ceranae*. Target sequences from the γGCS, GS, GPx-1, GPx-2, TrxR, and TPx genes were designed and amplified from the cDNA of *N. ceranae* (Fig. 4A; Fig. S1). Using the Bee Microbiome Toolkit, we assembled plasmids with an inverted arrangement of two promoters (pBTK150, pBTK151) and other previously designed parts for the production of dsRNA^24, 37^. We built six complete dsRNA-producing plasmids targeting different genes and transformed these plasmids into *S. alvi* M0351 by conjugation. We inoculated newly emerged honeybees with ~10^5^ cells of *S. alvi* bearing different plasmids that expressed dsRNA corresponding to the GFP coding sequence (pDS-GFP) or those expressed target sequences. Then, the bees were challenged by oral feeding with *N. ceranae* spores (10^4^ spores/bee), and 10 days later, we tested whether the *Snodgrassella*-produced dsRNA could inhibit the proliferation of *N. ceranae*.

**Fig. 4.**
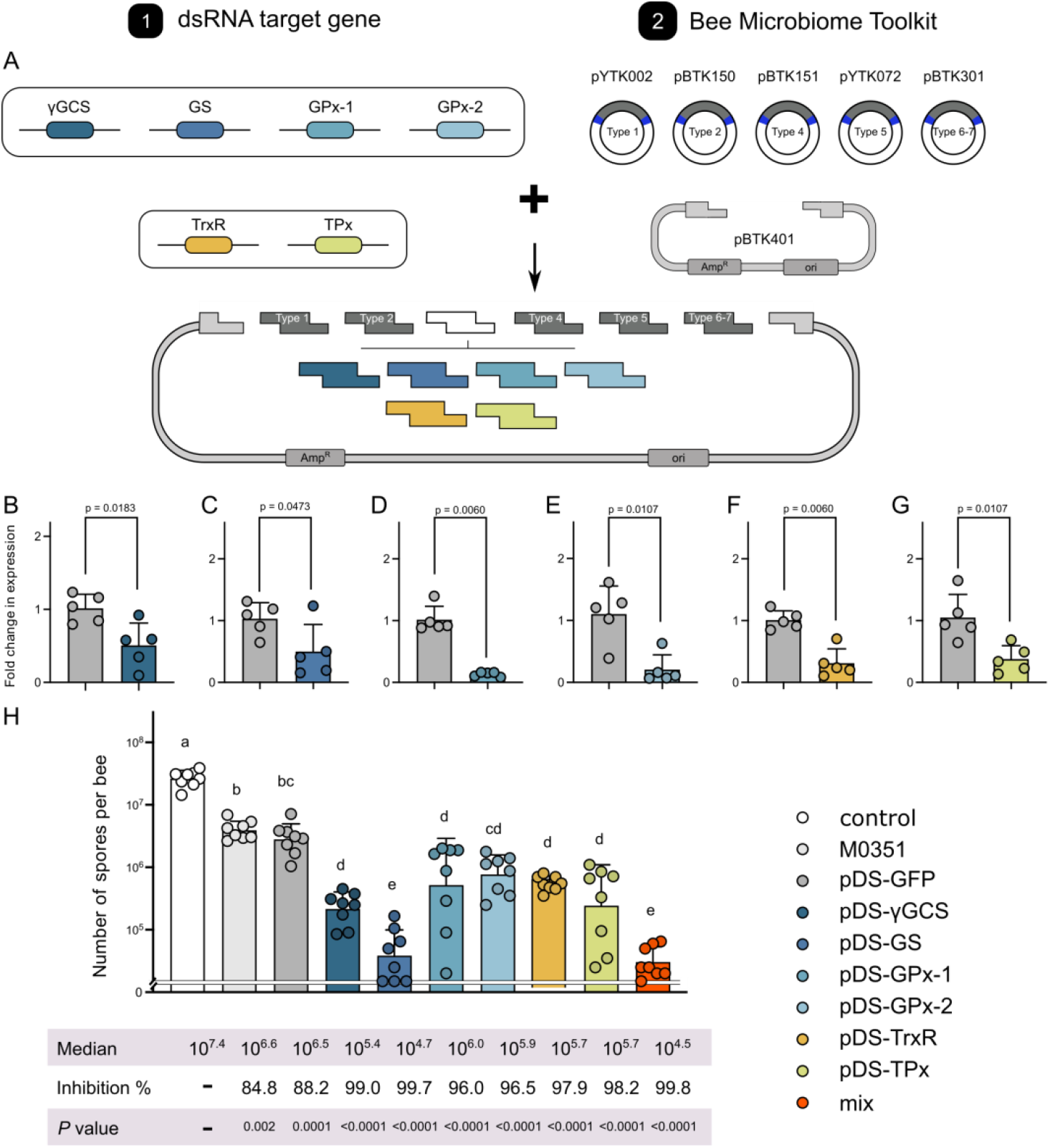
Recombinant *S. alvi* M0351 strains engineered to deliver dsRNA inhibit the infection of *N. ceranae*. (A) Schematic illustration of the Golden Gate assembly strategy for constructing plasmids expressing targeted dsRNA. (B-G) Engineered *S. alvi* strains to produce dsRNA targeting genes of the thioredoxin and glutathione systems inhibited the gene expression of *N. ceranae* in the midgut. (H) Inhibition of *N. ceranae* infection by engineered *S. alvi* M0351 strains. Each dot represents the spores number of an individual bee in the midgut. Letters above each bar stand for statistical differences between sampling sites (least-significant difference (LSD) test, *P* < 0.05). The median number of spores of each treatment group and the percent inhibition (inhibition %) of spore load relative to the non-colonized control are indicated.

We first extracted the RNA of *N. cerana* to confirm the depression of targeted pathways. Compared to the pDS-GFP off-target control, a significantly lower expression of target genes from *N. cerana* was identified (Fig 4B–G). Expression of all targeted genes is decreased by 50–86% in *N. ceranae* with different recombinant strains, suggesting that the dsRNA is delivered from the engineered *S. alvi* to allow diffusion to the parasite. After 10 days of dsRNA silencing, we evaluated the inhibitory capacity of various engineered *S. alvi* strains by quantifying the spore load with microscopic observation. First, both the wild-type *S. alvi* and the pDS-GFP provided protection compared with the controls without symbiont inoculation (Fig. 4H), confirming the role of *S. alvi* in defending against *N. ceranae*. Engineered *S. alvi* strains expressing γGCS, GPx-1, GPx-2, TrxR, or TPx dsRNA had a 99%, 96%, 96.5%, 97.9%, and 98.2% decrease in *N. ceranae* spore load, respectively. Notably, pDS-GS targeting the glutathione synthetase of the glutathione system showed the highest inhibition (99.7%) of the microsporidia spore invasion. Moreover, we also evaluated the effect of mixing engineered strains, and the inhibitory effect by a mixture of bacteria delivering all six dsRNA was better than the colonization with single strains.

## Discussion

*Nosema ceranae* is a microsporidian parasite initially identified from the *Apis cerana* in the 1990s^38^ and later detected in different honeybee species worldwide, becoming a globally distributed pathogen^27^. In this study, we found that *N. ceranae* employs the thioredoxin and glutathione system to relieve oxidative stress from the host for the adaptation in the midgut epithelium (Fig. 5). We showed that the core gut bacteria, *S. alvi*, triggers the redox response involved in honeybee gut immunity, which inhibits the proliferation of *N. ceranae* by up to 85.5%. Moreover, we successfully constructed engineered *S. alvi* M0351 based on the Bee Microbiome Toolkit and the Functional Genomics Using Engineered Symbionts procedure (FUGUES) to continuously produce dsRNA for critical genes of the *N. ceranae* thioredoxin and glutathione systems. Engineered *S. alvi* can stably re-colonize bees and repress the parasite’s thioredoxin and glutathione system-related gene expression. The γGCS and GS of the glutathione system are the most effective targeted genes for *N. ceranae* inhibition. Furthermore, the inhibitory effect by the mixture of all six engineered *S. alvi* strains showed the highest inhibition (99.8%).

**Fig. 5.**
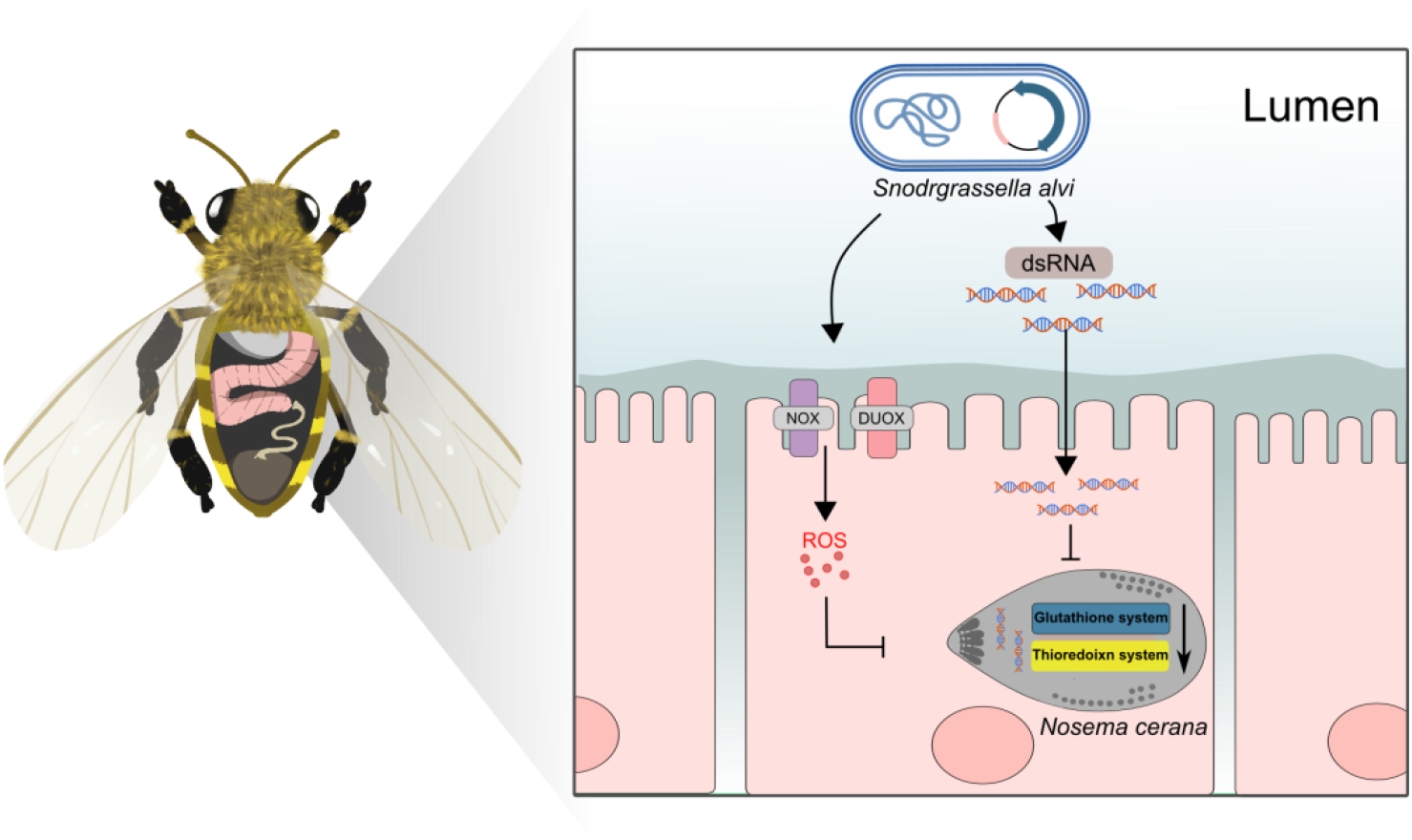
Graphical summary of the *N. ceranae* inhibition by the engineered *S. alvi* M0351 strains. By triggering *Duox* and *Nox* genes, *S. alvi* induces ROS production in the midgut of honeybees, which may kill *N. ceranae* via perturbation of redox homeostasis. *N. ceranae* may employ the thioredoxin and glutathione antioxidant systems to relieve the intense oxidative stress for intracellular proliferation. Engineered *S. alvi* M0351 expressing the dsRNA corresponding to the genes of the oxidation-reduction systems of *N. ceranae* can significantly inhibit parasitism in the honeybee gut.

The microsporidia *N. ceranae* is an obligate intracellular parasite that develops in the ventricular epithelia of *A. mellifera*, and the spores can spread the infection quickly across epithelial cells^39^. The concept that the immune activation of honey bees plays a role against *N. ceranae* is not new, but the exact mechanism remains unclear. In all invertebrates, including insects, the adaptive system is missing, and hence, defense is entirely ensured by the innate immune system. The innate immune system provides organisms with a rapid, non-specific first line of defense against colonization by pathogenic microorganisms^40^. Honeybees’ cell-mediated innate immune system consists of hemocytes, which produce ROS essential for cell signaling and pathogen clearance^41^. ROS are produced during the recognition and phagocytosis response against the foreign bodies inside the cells^42^. In insects, ROS was shown to be particularly important in the fight against parasites and pathogenic bacteria in the gut^27, 43^. The production of ROS has been observed following the infection of *Aedes aegypti* mosquito by different microsporidian species^44^. In honeybees, genes involved in the production of ROS are overrepresented in the midgut upon the spore infection, suggesting that the increased ROS level is an immune response against intracellular parasitism^19^. Our results confirmed that the ROS-associated immune response in the epithelial cells is indispensable for preventing microsporidia infection in bees.

Two ROS-producing enzymes, the NADPH oxidases Duox and Nox, are identified in *A. mellifera*, similar to *Drosophila*. Here, we found that the expression of both *Duox* and *Nox* genes was activated in the midgut of bees post-colonization by *S. alvi*. It has been documented that *S. alvi* protects honeybees from the opportunistic pathogen *Serratia marcescens* by triggering the Toll and Imd pathway to upregulate the expression of host antimicrobial peptides abaecin, apidaecin, and hymenoptaecin^45^. *S. alvi* colonizes the honeybee gut in contact with the gut epithelia and forms a dense biofilm, which may stimulate pattern recognition receptors such as Toll-like receptors of bees^45^. Interestingly, the activation of Toll-like receptors possibly conjugates the NADPH oxidases, which are involved in the ROS production on the membranes of the endosome of cells^46^. Notably, bees mono-colonized with *S. alvi* strain inhibit the proliferation of *N. ceranae* by up to 85.5%. In contrast, bees colonized by other gut members did not show a significant reduction of *N. cerana*. Thus, the ROS immune response may be activated by *S. alvi* by regulating the Toll signaling and the NADPH complex. The ROS system is also a gut-immune immune response involved in gut-microbe homeostasis^18^. In honeybees, the *N. ceranae* infection can perturb the gut composition, and a normal microbiota is required for host resistance to *N. ceranae*^47, 48^. The bore gut members of honeybees, including the *S. alvi* and *Lactobacillus apis*, have been shown to activate the humoral innate immune system to produce antimicrobial peptide (AMP), which protects against pathogens^22, 23^. Our data shows that the bee gut bacteria also plays a pivotal role in controlling microsporidia invasion by inducing de novo generation of ROS^18^.

To protect itself against host defenses of ROS, the parasite may have an internal antioxidant system to maintain a normal redox state. For example, *Plasmodium falciparum* scavenges ROS from hosts by employing antioxidant systems, including the NADPH-dependent thioredoxin and the glutathione system ^34, 49^. We found that *N. ceranae* also possesses balanced redox pathways, including the thioredoxin and glutathione systems, which may be necessary for counteracting ROS attacks from the host. Analyzing the time series of gene expression of *N. ceranae* colonizing the bee gut, we found significant upregulation in the expression of thioredoxin and glutathione system genes, suggesting that the maintenance of the normal redox state is significant in the invasion process of *N. ceranae*. Previous studies that used RNAi targeting on variable *N. ceranae* genes, such as the polar tube protein 3 (ptp3) and the spore wall protein (SWP), reduce parasite load and improve the physiological performance of honeybees^50, 51^. Our results showed that the delivery of dsRNA corresponding to γGCS and TrxR led to a significant reduction in spore load by 82% and 85%, respectively, indicating that the endogenous antioxidant enzymes of *N. ceranae* provide a novel therapeutic target for the control of the parasitic invasion in honeybees.

Although RNAi is widely used, it is expensive for application in agricultural fields, and its efficiency is also unsatisfactory. Recently, researchers have engineered host-associated bacterium to produce dsRNA as a novel delivery modality. Taracena *et al*. used engineered *Escherichia coli* to produce dsRNA to control the *Rhodnius prolixus* parasite, a vector of the Chagas disease^52^. Although laboratory *E. coli* is easy to be engineered, the symbiotic bacteria with minimal fitness cost and stable association with the host are more promising for *in vivo* treatment ^53^. Recently, Leonard *et al*. have designed the Functional Genomics Using Engineered Symbionts (FUGUES) procedure for engineering the native bacterial species of honeybees^25^. The engineered *S. alvi* produces dsRNA to inhibit parasitic *Varroa* by inducing mite RNAi response^37^. It documents that the RNA produced by the recombinant strains can be transported to the gut, hemolymph, and head of bees, which alters host physiology and behavior. *N. ceranae* infects and proliferates intracellularly in the epithelial cells of the midgut^38^. While *Snodgrassella* mainly localizes to the ileum region of the hindgut, it also colonizes the midgut intensively^35^. Our results illustrated that the recombinant *S. alvi* strain stably re-colonizes both ileum and midgut post-inoculation, and the plasmid exhibit a robust expression *in vivo*. Thus, the dsRNA produced by the symbiotic *S. alvi* may enter into the midgut cells and destroy the redox homeostasis of *N. ceranae*. Here, we targeted all six genes involved in the glutathione and thioredoxin systems of the microsporidia. Interestingly, the repression of the GS gene showed the highest inhibition efficiency. GS is not subject to feedback inhibition by GSH and is important in determining overall GSH synthetic capacity, specifically under pathological conditions^54^. In addition, the inhibition of a mixture of the recombinant strains is more significant than the single-targeting strains, suggesting a more effective perturbation of the parasitic antioxidant system. *S. alvi* and other bee gut bacteria can be naturally transmitted within the colony via social contact, and engineered *S. alvi* strains are transferred between co-housed bees, which suggests that the use of native gut bacteria can facilitate the treatment of individual bees from an entire colony^37^. Although gene escape is a major ethical issue for the application of engineered bacteria, honeybee symbiotic bacteria are generally restricted to the bee gut environment^21^. Moreover, *S. alvi* shows even more strict host specificity that only specific strains are associated with different bee species or even individuals within the colony^55^. Thus, symbiont-mediated RNAi provides a new tool to improve resilience against current and future challenges to honey bee health.

## Materials and Methods

### Generation of microbiota-free and mono-colonized honeybees

Honeybees (*A. mellifera*) used in this study were from colonies maintained in the experimental apiary of the China Agricultural University. Pupae and newly emerged bees used in all the experiments were obtained from brood frames taken from the experimental hives and kept in an incubator at 35 °C, with a humidity of 50%. All honeybee gut bacterial strains used in this study are listed in Table S1. *Snodgrassella alvi* M0351, *Bifidobacterium choladohabitans* W8113, and *Gilliamella apicola* B14384H2 isolated from honeybee guts were grown on HIA supplemented with 5% (vol/vol) sterile sheep blood (Solarbio). *Bombilactobacillus mellis* W8089 and *Lactobacillus apis* W8172 were grown on MRS agar plates (Solarbio) supplemented with 0.1% L-cystine and 2.0% fructose.

Microbiota-free (MF) bees were obtained as described by Zheng *et al*.^56^. In brief, we manually removed pupae from brood frames and placed them in sterile plastic bins. Newly emerged MF bees were kept in axenic cup cages with sterile sucrose syrup for 24h. For each mono-colonization setup, 20–25 MF bees were placed in a cup cage and fed bacterial culture solutions for 24 hours. Colonization levels were determined by colony-forming units from dissected guts, as described by Kwong *et al*.^56^ 1 mL of 1×PBS was combined with 1 mL of sucrose solution and 0.3 g of sterilized pollen for the MF group. For the mono-colonization bees, glycerol stock of bee gut strains was resuspended in 1 mL 1×PBS at a final concentration of ~10^8^ CFU/mL and then mixed with 1 mL sterilized sucrose solution with 0.3 g of sterilized pollen. The bees were incubated at 35°C and RH 50% until Day 7.

### *N. ceranae* spore purification

*N. ceranae* spores were isolated from worker honeybees collected from heavily infected colonies in the summer of 2022. After immobilizing bees by chilling them on ice, the guts were removed from individual bees with forceps. The midguts of infected honeybees were homogenized in distilled water and filtered using Whatman filtering paper. The filtered suspension was centrifuged at 3,000×g for 5 minutes, and the supernatant was discarded. The re-suspended pellet was purified on a discontinuous Percoll (Sigma-Aldrich, St. Louis, MO) gradient of 5 ml each of 25%, 50%, 75%, and 100% Percoll solution. The spore suspension was overlaid onto the gradient and centrifuged at 8,000 × g for 10 minutes at 4°C. The supernatant was discarded, and the spore pellet was washed by centrifugation and suspension in distilled sterile water^57^. The number of spores was quantified using a Fuchs-Rosenthal hemocytometer. The identity of the isolated *N. ceranae* or *N. apis* was determined by amplifying the ribosomal RNA gene sequences with species-specific primers (Table S3)^58^.

### Bees mono-colonized with gut symbionts challenged with *N. ceranae*

To accurately control the number of *N. ceranae* cells infecting each bee individual, bees were orally fed the same amount of *N. ceranae* spores. Each bee was starved for 2 hours and given 2 μl of a 50% sucrose solution containing 10^4^ *N. ceranae* spores. After 10 days, the number of spores in the intestinal specimen of infected bees was quantified as described by Huang *et al*.^47^. The midguts were dissected, resuspended in 500 μl of double-distilled water, and then subjected to vortex mixing. The suspension was put onto the hemocytometer for microscopic observation.

### Honeybee gut RNA extraction and quantitative PCR

Each dissected gut was homogenized with a plastic pestle, and total RNA was extracted from individual samples using a Zymo Quick-RNA Tissue/Insect Microprep Kit (Zymo Research, #R2030). RNA was eluted into 50 μL of RNase-free water and stored at −80 °C prior to reverse transcription. cDNA was synthesized using the HiScript III All-in-one RT SuperMix Perfect for qPCR (Vazyme). Quantitative real-time PCR was performed using the ChamQ Universal SYBR qPCR Master Mix (Vazyme) and QuantStudio 1 Real-Time PCR Instrument (Thermo Fisher Scientific, Waltham, MA, USA) in a standard 96-well block (20-μl reactions; incubation at 95 °C for 3 min, 40 cycles of denaturation at 95 °C for 10 s, annealing/extension at 60 °C for 20 s). The primers for the genes of *Duox* (LOC551970) and *Nox* (LOC408451) of *A. mellifera* were designed with IDT qPCR PrimerQuest Tool (https://www.idtdna.com/pages/tools/primerquest) (Table S3). The actin gene of *A. mellifera* was used as the control, and the relative expression was calculated using the 2^−ΔΔCT^ method ^59^.

### *In vivo* detection of reactive oxygen species

Three days after inoculation, the midguts of the honeybees mono-colonized with *S. alvi* M0351 and the MF bees were dissected in PBS containing 50 μM intracellular ROS-sensitive fluorescent dye dihydroethidium (Invitrogen, #C10422). The tubes were placed in the dark for 10 min at room temperature. Then, the midguts were washed twice with a fresh dye-free PBS, and the tissues were immediately transferred to an μ-Dish^35 mm, high^ microscope dishes (Ibidi, #81156). We imaged the gut tissues on a Zeiss 910 Laser Scanning Confocal microscope with a 20× objective.

### Measurement of the H_2_O_2_ production

The generation of H_2_O_2_ was determined using the Hydrogen Peroxide Assay Kit (Beyotime Biotech). In this assay, H_2_O_2_ converts Fe^2+^ to Fe^3+^, which then reacts with xylenol orange dye to become purple with a maximum absorbance at 560 nm. The midguts were homogenized in 200 μL lysis buffer and centrifuged at 12,000 × g at 4 °C for 5 min, and the supernatant was collected. Aliquots of 50 μL of supernatants and 100 μL of test solutions from the Hydrogen Peroxide Assay Kit were incubated at room temperature for 20 minutes and measured immediately with a spectrometer at a wavelength of 560 nm. The measurement was repeated three times for each sample.

### RNA isolation of *N. ceranae*

To extract the RNA of *N. ceranae*, the honeybee gut was individually transferred into 2 ml tubes. Each tube contained 100 μL sterile 1.4-mm zirconium silicate grinding beads (Quackenbush). One milliliter of TRIzol reagent (Ambion) was added to the tube, disrupting the samples using the FastPrep. The samples were treated with DNase I (Invitrogen) to remove genomic DNA contamination. The purity and quantity of RNA samples were determined using a NanoDrop 8000 spectrophotometer (Thermo Fisher Scientific). cDNA was synthesized using the HiScript III All-in-one RT SuperMix Perfect for qPCR (Vazyme) and stored at −20°C.

### *Nosema* inoculation and Nanocarrier-mediated dsRNA feeding assay

To produce the double-stranded RNA of the γGCS (AAJ76_1100057370) and TrxR (AAJ76_5800012528) genes, the coding regions of the genes were amplified from *N. ceranae* cDNA with forward and reverse primers containing the T7 promoter sequence at their 5′ends (5′-TAATACGACTCACTATAGGGCGA-3′). The partially amplified segments of the genes were cloned into the pCE2-TA-Blunt-Zero vector (Vazyme, China) and verified by Sanger sequencing. The fragment was amplified from the plasmid using specific primers with a T7 promoter and then used for dsRNA synthesis using the T7 RNAi Transcription Kit (Vazyme, China). The fragment amplified from the GFP gene (MH423581.1) was used as the control. The sequences of the primers are given in Table S3. Here, we used the star polycation as a gene nanocarrier to protect dsRNA molecules from enzymatic degradation and promote their translocation across cell membranes^60^. The nanocarrier was mixed with γGCS and TrxR dsRNA gently at a mass ratio of 1:1. (The final concentration for both SPc and dsRNA was 100 ng/μL.) The final concentrations for dsRNA + nanocarrier, and sucrose were 100 ng/μl and 50% (wt/vol), respectively. Newly emerged bees were removed from the frames and kept without food for at least 2 h before the subsequent *N. ceranae* inoculation. Individual bees were fed 2 μL of spores suspensions prepared by mixing purified spores into 50% sucrose (~10^4^ spores/μL). From the day after *N. ceranae* inoculation, honeybees from each treatment were fed on different dsRNA mixtures in an incubator at 35°C The dsRNA mixture was supplied daily, and each bee ingested about 10 μg of dsRNA per day.

The treatment effect of dsRNA was determined by comparing the spore production rate for individual honey bees. The *N. ceranae* spore production rate was measured by counting the spores from the extracted midgut of live honey bees 15 days after inoculation. To investigate the effect of dsRNA treatment on the expression of each target gene of *Nosema*, qRT-PCR was performed after 15 days of dsRNA treatment. After extracting the midguts from honeybees treated with *Nosema* and dsRNA, the total RNA was extracted. cDNA was synthesized using the HiScript III All-in-one RT SuperMix Perfect for qPCR (Vazyme). Each gene-specific primer is given in Table S3. The β-tubulin gene of the *N. ceranae* was used as the control, and relative expression was analyzed using the 2^−ΔΔCT^ method ^59^.

### Vector construction to express dsRNA expression and *S. alvi* M0351 egineering

All the plasmids and MFD*pir* were kindly donated by the Moran Lab and Barrick Lab (University of Texas at Austin). We designed dsRNA-producing plasmid parts based on the previously published Bee Microbiome Toolkit and functional genomics using engineered symbionts procedure (FUGUES) (Fig. 4A)^25^. First, PCR is used to amplify the knockdown region γGCS, GS, GPx-1, GPx-2, TrxR, and TPx from the cDNA of *N. ceranae* and append BsaI cut sites to each end. Following PCR, amplicons are purified and cloned into a dsRNA expression vector. We combined previously designed parts pYTK002 (Type 1), pBTK150(Type 2), pBTK151(Type 4), pYTK072 (Type 5), pBTK301 (Type 6-7), and pBTK401 (Type 8) (Addgene_65109, Addgene_183127, Addgene_65179, Addgene_183126, Addgene_110593, Addgene_110597), and dsRNA target sequence (Type 3) to assemble complete plasmids that express dsRNA of the target sequence^37^. Golden Gate assembly reactions were performed as previously described^24^, and enzyme BsaI-HFv2 (New England Biolabs) was used to increase assembly efficiency.

Assemblies were transformed into electroporated into *E. coli* donor strain MFD*pir*. The plasmids were verified with Sanger sequencing. MFD*pir* cells with the dsRNA expression vector and *S. alvi* cells are grown, washed, and combined to initiate conjugation. Then, this mixture is plated on media containing 0.30 mM DAP. The next day, cells are scraped, washed, and plated on media containing 100 μg/mL ampicillin but without DAP to select for *S. alvi* cells that have acquired the plasmid and against MFD*pir* cells. Transconjugant *S. alvi* colonies are passaged onto a second plate containing 100 μg/mL ampicillin. These transconjugants can be confirmed to be pure *S. alvi* cultures by performing 16S rRNA sequencing to ensure no unexpected contaminants have been introduced during the conjugation process.

After 2–3 d of growth, we scraped the engineered *S. alvi* grown on the plates into PBS. These cells were spun in a centrifuge (3824 × g, 5 min) and then resuspended in 500 μL PBS. Engineered *S. alvi* was diluted in 500 μL 1×PBS at a final concentration of ~10^8^ CFU/mL and combined with 500 μL of a 1:1 sucrose: water solution supplemented with 200 μg/mL ampicillin. We fed engineered *S. alvi* solutions to age-controlled newly emerged worker bees for 24 h (pDS-γGCS, pDS-GS, pDS-GPx-1, pDS-GPx-2, pDS-TrxR, pDS-TPx) and non-targeted (pDS-GFP) served as a negative control group. The next day, each bee was given 2 μl of a 50% sucrose solution containing 10^4^ *N. ceranae* spores. After ten days, honeybee gut was collected to quantify the number of *N. ceranae* spores, and gene knockdown was validated using qPCR on the cDNA of *N. ceranae* synthesized as described above.

To test whether engineered *S. alvi* robustly colonizes bees, we inoculated bees with *S. alvi* transformed with a plasmid expressing GFP. Firstly, we transformed strain M0351 with a stable plasmid pBTK519 expressing GFP from the Bee Microbiome Toolkit platform ^24^ and inoculated bees with *S. alvi* M0351::pBTK519 (~10^5^cfu/bee). After every five days, we dissected bees, homogenized their whole guts in 500 μL PBS, and plated dilutions onto HIA plates with a final concentration of 100 μg/mL ampicillin to estimate CFUs of *S. alvi* in the gut. The number of fluorescent and non-fluorescent colonies on the plates was quantified to track the stability of engineered strains over time. After 15 days, we dissected the guts and imaged them on a Laser Scanning Confocal microscope.

## Acknowledgments

We thank the Moran Lab and Barrick Lab (University of Texas at Austin) for donating plasmids. We thank Elijah Powell, Jeffrey Barrick, and Nancy Moran for their suggestions in gut bacteria engineering. We thank Jie Shen and Shuo Yan (China Agricultural University) for their assistance in the nanoparticle RNA delivery system. This work was supported by National Key R&D Program of China (Grant No. 2019YFA0906500), National Natural Science Foundation of China Project 32170495 and 31760715.

## Supplementary Information

**Fig. S1.**
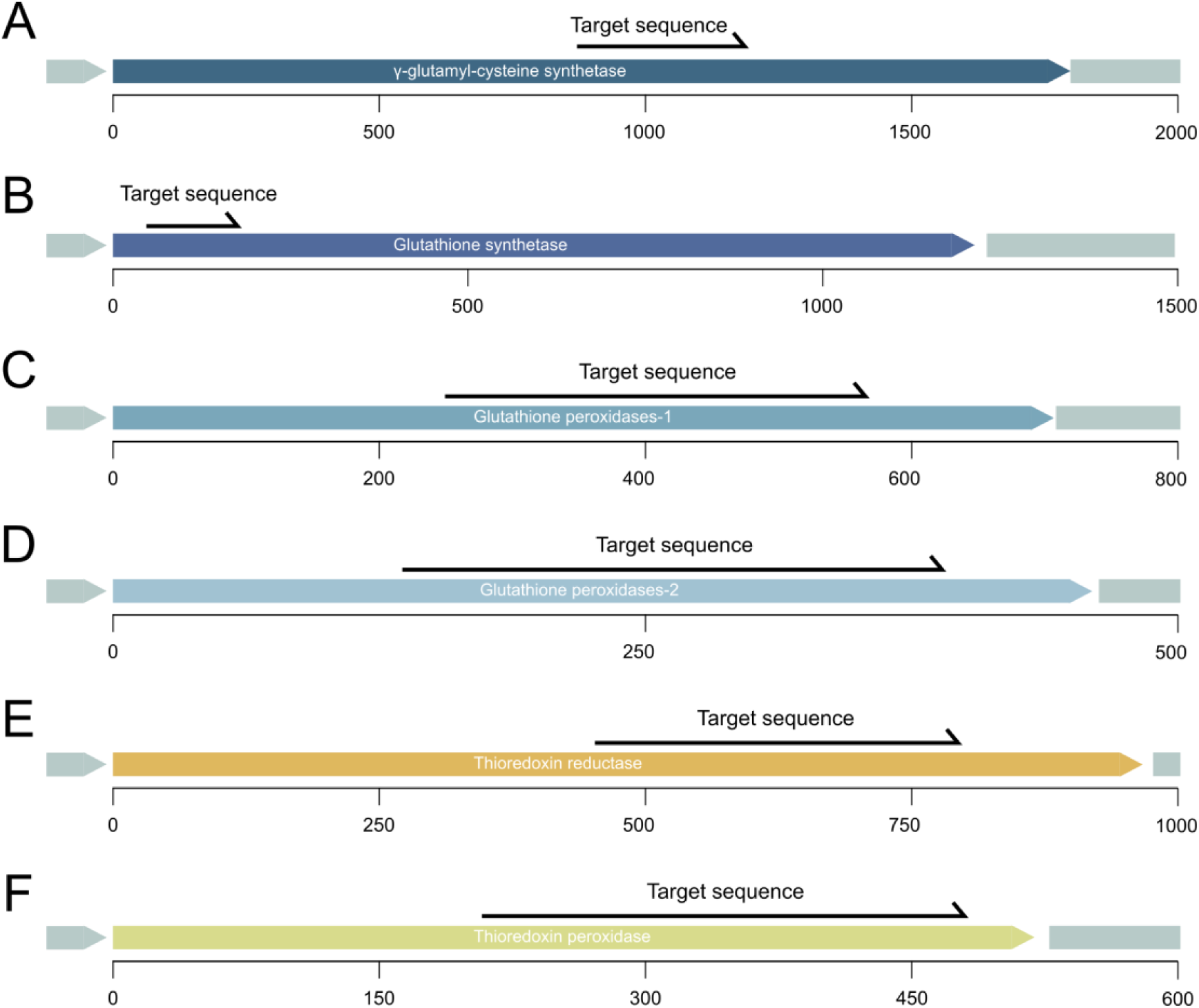
Design of *N. ceranae* targets. This diagram shows the overall gene organization for *N. ceranae* γ-glutamyl-cysteine synthetase (A), glutathione synthetase (B), Glutathione peroxidases-1 (C), Glutathione peroxidases-2 (D), thioredoxin reductase (E), and thioredoxin peroxidase (F). The targeted region in each gene is denoted, and the designed target sequences are listed in Table S4.

**Table S1.** List of bacterial strains.

| Species and strain | Source |
| --- | --- |
| *E. coli* MFDpir | <sup>24</sup> |
| *E. coli* DH5α | Vazyme |
| *Snodgrassella alvi* M0351 | This study |
| *Bifidobacterium choladohabitans* W8113 | This study |
| *Bombilactobacillus mellis* W8089 | This study |
| *Lactobacillus apis* W8172 | This study |
| *Gilliamella apicola* B14384H2 | This study |
| *Snodgrassella alvi* M0351 | This study |

**Table S2.** Plasmid list.

| Name | Use | Source |
| --- | --- | --- |
| pBTK519 | Constitutive GFP | 24 |
| pYTK002 | Type 2 YTK/BTK connector sequence part plasmid | 24 |
| pYTK072 | Type 5 YTK/BTK connector sequence part plasmid | 24 |
| pBTK301 | Type 6-7 BTK bridge connector sequence part plasmid | 24 |
| pBTK401 | Type 8 origin of replication and origin of transfer plasmid, *rsf1010* broad-host-range origin | 24 |
| pBTK150 | Type 2 BTK part: terminator, CP25 promoter, no RBS | 37 |
| pBTK151 | Type 4 BTK part: reverse CP25 promoter, terminator, no RBS | 37 |
| pYTK001_T1T2 | Insulated part vector with flanking terminators | 37 |
| pDS-GFP | Control dsRNA GFP | 37 |
| pDS-γGCS | dsRNA target γGCS | This Study |
| pDS-GS | dsRNA target GS | This Study |
| pDS-GPx-1 | dsRNA target GPx-1 | This Study |
| pDS-GPx-2 | dsRNA target GPx-2 | This Study |
| pDS-TrxR | dsRNA target TrxR | This Study |
| pDS-TPx | dsRNA target TPx | This Study |

**Table S3.**
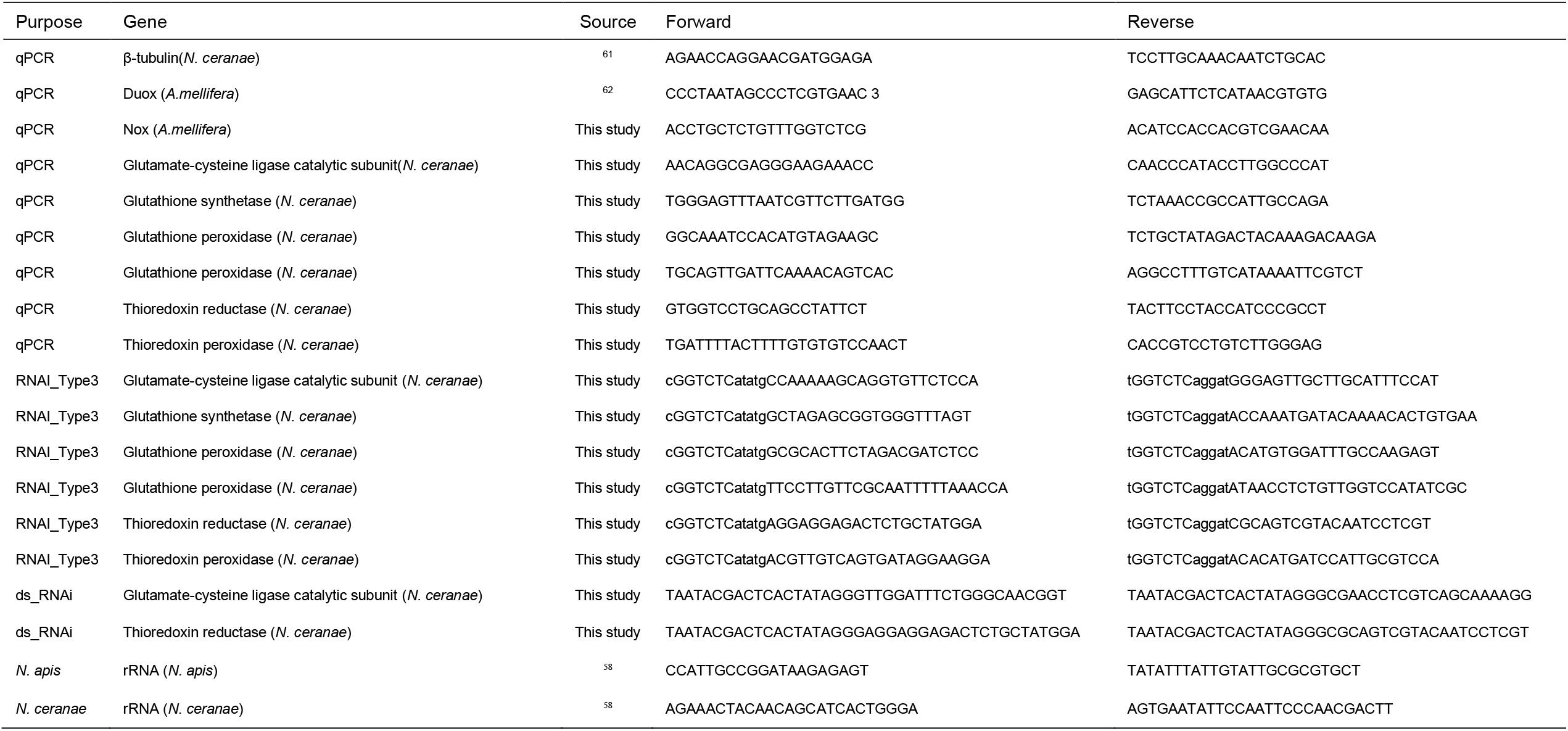
Primer list.

**Table S4.**
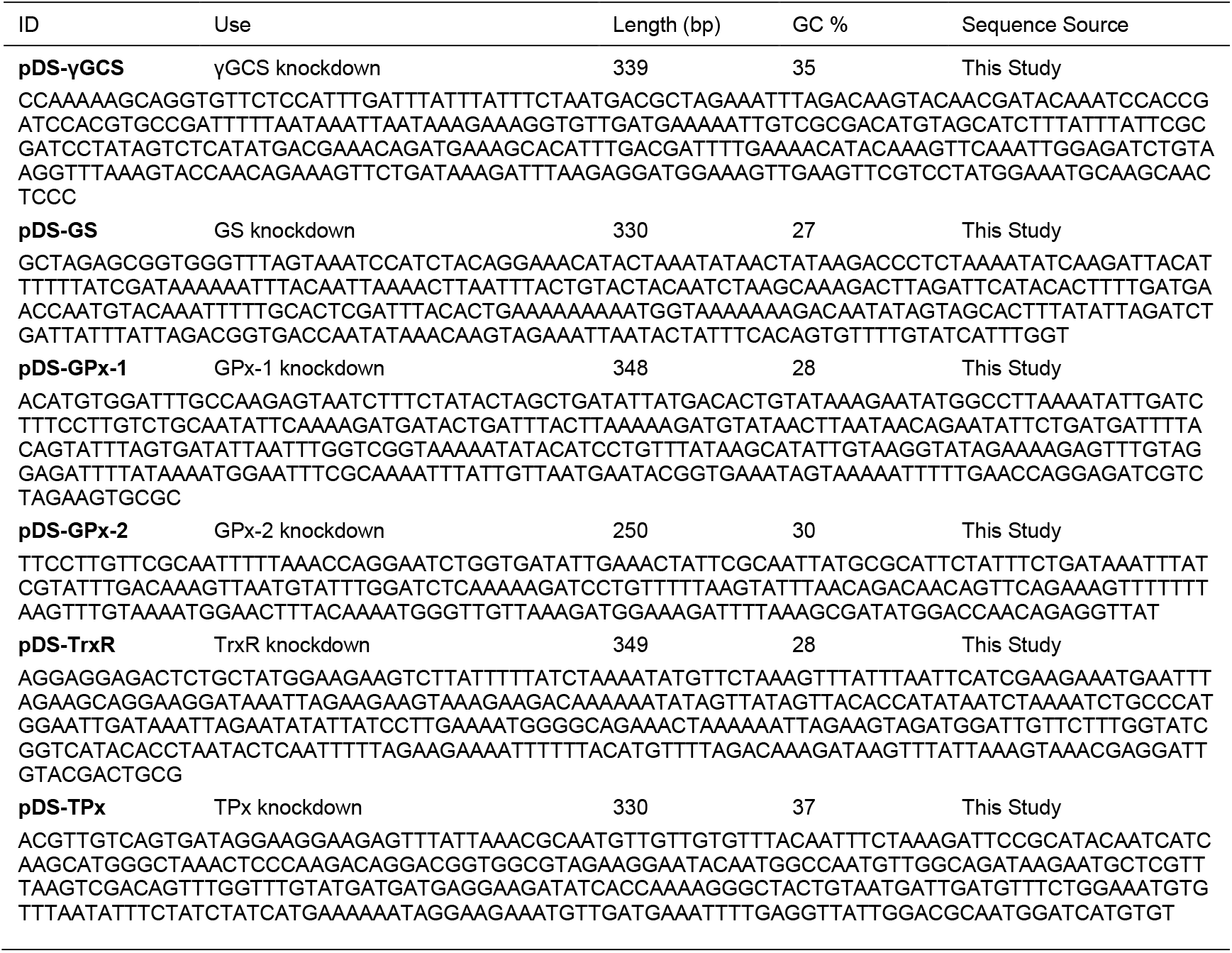
Target sequences of dsRNA.

